# Transcriptomic profiling of canine atrial fibrillation models after one week of sustained arrhythmia

**DOI:** 10.1101/2021.02.04.429512

**Authors:** Francis J.A. Leblanc, Faezeh Vahdati Hassani, Laura Liesinger, Xiaoyan Qi, Patrice Naud, Ruth Birner-Gruenberger, Guillaume Lettre, Stanley Nattel

## Abstract

**Background:** Atrial fibrillation (AF), the most common sustained arrhythmia, is associated with increased morbidity, mortality, and health-care costs. AF develops over many years and is often related to substantial atrial structural and electrophysiological remodeling. AF may lack symptoms at onset and atrial biopsy samples are generally obtained in subjects with advanced disease, so it is difficult to study earlier-stage pathophysiology in humans.

**Methods:** Here, we characterized comprehensively the transcriptomic (miRNAseq and mRNAseq) changes in the left atria of two robust canine AF-models after one week of electrically-maintained AF, without or with ventricular rate-control via atrioventricular node-ablation/ventricular pacing.

**Results:** Our RNA-sequencing experiments identified thousands of genes that are differentially expressed, including a majority that have never before been implicated in AF. Gene-set enrichment analyses highlighted known (e.g. extracellular matrix structure organization) but also many novel pathways (e.g. muscle structure development, striated muscle cell differentiation) that may play a role in tissue remodeling and/or cellular transdifferentiation. Of interest, we found dysregulation of a cluster of non-coding RNAs, including many microRNAs but also the *MEG3* long non-coding RNA orthologue, located in the syntenic region of the imprinted human *DLK1-DIO3* locus. Interestingly (in the light of other recent observations), our analysis identified gene-targets of differentially expressed microRNAs at the *DLK1-DIO3* locus implicating glutamate signaling in AF pathophysiology.

**Conclusions:** Our results capture molecular events that occur at an early stage of disease development using well-characterized animal models, and may therefore inform future studies that aim to further dissect the causes of AF in humans.

## INTRODUCTION

Atrial fibrillation (AF) is the most common sustained arrhythmia, with an estimated lifetime risk of 22%-26% and association with increased morbidity and mortality^1^. Despite advances in antiarrhythmic therapies, their suboptimal efficacy and adverse effects have limited their use^2^. Therefore, there is a need to further characterize fundamental arrhythmia mechanisms in order to discover new therapeutic targets^2^. Although AF is known to be a final common endpoint of atrial remodeling resulting from a variety of heart diseases, it can also be, in turn, a cause of remodeling. This vicious cycle is called “AF begets AF”^3^ and explains the progressive nature of this arrhythmia and the complexity of its management.

Atrial remodeling is characterized by ion channel dysfunction, Ca^2+^ handling abnormalities, and structural changes, which result in AF induction and maintenance^4, 5^. Heart disease, and even rapid atrial activity itself, cause the development of atrial fibrosis, which is a hallmark of structural remodeling. The degree of fibrosis is positively correlated with the persistence of AF^6^. Atrial cardiomyocytes subjected to rapid activation release factors that induce fibroblast-to-myofibroblast differentiation that leads to increased collagen synthesis^7^.

Any arrhythmia causing a rapid ventricular rate, including AF, is a well-recognized inducer of ventricular dysfunction, so-called “arrhythmia-induced cardiomyopathy”^8^. Heart failure enhances atrial stretch and sympathetic tone, making AF more resistant to rate- or rhythm-control treatments^9^. AF promotion results from the rapid atrial rate, but rapid ventricular rates due to inadequate rate-control also promote AF-related atrial remodeling with a different profile from the remodeling produced by rapid atrial rate alone^10^. Radiofrequency atrioventricular node ablation (AVB) with right ventricular pacing is a nonpharmacological strategy for rate control that can improve symptoms and outcomes^11^.

Our previous work in canine AF models showed that maintaining AF for one week by rapid atrial pacing activates fibroblasts, collagen gene expression and cardiomyocyte ion channel changes, without yet causing fibrosis^12^. Continued electrical maintenance of AF for 3 weeks produces fibrosis, but electrically-maintained AF with ventricular rate control through AVB produces less profibrillatory remodeling than 3 weeks of AF alone^10^. However, how AF with and without AVB impact atrial remodeling at the molecular level has not yet been assessed comprehensively. To answer this question, we took advantage of our well-characterized AF dog models and performed RNA-sequencing (RNA-seq) of cardiomyocyte-enriched atrial samples after one week to capture the molecular actors of atrial remodeling. In comparison with control (CTL) dogs, we found thousands of mRNAs, long non-coding RNA (lncRNA) and microRNA (miRNA) that are differentially expressed (DE) in the atria of the canine AF models. Pathway analyses of the transcriptomic data highlighted known biological processes, but also potential novel modulators of arrhythmia initiation which may shed new light on our understanding of AF in humans.

## METHODS

All results and R code are available at https://github.com/lebf3/DogAF.

### Canine atrial fibrillation model

A total of 18 adult mongrel dogs of either sex, weighing 18-32 kg, were obtained from LAKA Inc and randomly assigned to control (CTL) group (n=6) and two canine AF-models (n=6/group) (**Supplemental Table I**). We selected 6 animals per group based on previously published RNAseq studies and our previous experiments with these models. Further, the number of DE genes identified in our analyses (post hoc) indicates that this sample size is sufficient to capture the main transcriptional changes that occur in the atrium of these dog models. Animals were handled in accordance with the “Guide for the Care and Use of Laboratory Animals” established by the National Institutes of Health as approved by the Montreal Heart Institute Ethics Committee (2016-47-01, 2019-47-03 for control dogs, 2015,47-01, 2018.47.12 for AF dogs).

To induce AF, animals were subjected to atrial tachypacing without (AF)^13^ and with (AF+AVB)^14^ atrioventricular-node ablation under 0.07 mg/kg acepromazine (IM), 5.3 mg/kg ketamine (IV), and 0.25 mg/kg diazepam (IV), and 1.5% isoflurane anesthesia. In the AF group, a bipolar pacing lead with fluoroscopic guidance was placed in the right atrial appendage (RAA). In the AF+AVB group, pacing leads were inserted into the RAA and right ventricular apex. Pacing leads were connected to a subcutaneous pacemaker implanted in the neck (right side). In the AF+AVB group, radiofrequency catheter ablation was used to create AF+AVB. For this purpose, a quadripolar catheter with fluoroscopic guidance was placed across the tricuspid valve via the right femoral vein. Radiofrequency energy was then used to perform ablation when action potential at the His bundle was detected. Twenty-four to seventy-two hours after surgery, dogs in the AF group were subjected to AF-maintaining atrial tachypacing at 600 bpm for seven days. In the AF+AVB group, RA and right ventricle were paced at 600 and 80 bpm, respectively. In animals of the CTL group, no pacemaker was inserted. No adverse event was recorded and no dog was excluded.

### Enrichment of dog atrial cardiomyocytes

Cardiomyocytes were enriched from the left atrium (LA) with enzymatic digestion through the coronary artery-perfused Langendorff system, as previously described^15^. Briefly, dogs were anesthetized with 2 mg/kg morphine (IV) and 120 mg/kg alpha-chloralose and mechanically ventilated. Hearts were aseptically and quickly removed after intra-atrial injection of 10,000 U heparin and placed in Tyrode’s solution containing 136 mM NaCl, 5.4 mM KCl, 2 mM CaCl2, 1 mM MgCl2, 10 mM dextrose, 5 mM HEPES, 0.33 NaH2PO4 (pH was adjusted to 7.3 with NaOH). The left coronary artery of the isolated heart was cannulated, and the LA was dissected free and perfused with 100% oxygenated Tyrode’s solution (37°C, 1.8 mM Ca^2+^). The arterial branches were ligated to have a leak-free system, and LA tissues were perfused with Ca^2+^-free Tyrode’s solution for ~10 minutes, followed by ~1-hour perfusion with ~0.45 mg/mL collagenase (CLSII, Worthington, Lakewood, NJ) and 0.1% bovine serum albumin (Sigma–Aldrich, Oakville, ON) in Ca^2+^-free Tyrode’s solution for enzyme digestion. Digested tissue was removed from the cannula and cut into small pieces, and atrial cardiomyocytes were harvested.

### RNA-seq/miRNA-seq

#### Library preparation and sequencing

mRNA and miRNA libraries were prepared at Genome Québec. mRNA libraries were made with the NEBNext_dual kit (rRNA-depleted stranded) and sequenced on NovaSeq 6000 S2 PE100 Illumina platform generating 32-123M Paired-end reads per sample. miRNA libraries were prepared with TruSeq smRNA and sequenced on the HiSeq 4000 SR50 Illumina platform generating 10-12M reads per sample.

#### Bioinformatic processing and DE analysis

The complete analysis can be found at https://github.com/lebf3/Dog_AF_transcriptomic. Briefly, mRNA reads were pseudomapped on reference transcriptome CanFam3.1.98 with Kallisto^16^ with the options quant -t 5 -b 100 and the rest as default. We aggregated transcripts by genes with tximport^17^ and quantified with DESeq2^18^ Genes with 0 reads in more than 12 samples were removed. Shrunken log_2_ transformed expression corrected for library size, with and without fibroblast fraction as a covariate (within DEseq2’s model) were then analyzed for DE with Wald test for all pairwise comparisons of CTL, AF and AF+AVB and likelihood ratio test for total assay DE. We did not adjust our differential gene expression analyses for biological sex because no genes where DE between female and male dogs in our experiment. We plotted the PCA with fibroblast fraction as a covariate from log_2_ transformed expression values corrected with Limma’s removeBatchEffect() function^19^ for visualization of fibroblast effect on the top 1000 most variable genes. We then compared sets of GENEIDs found to be up (L2FC > 0 & *p* < 0.01) or down (L2FC < 0 & *p* < 0.01) in all possible contrasts. For miRNAs, we trimmed reads using fastp^20^ with default settings and aligned them to CanFam3.1.98 genome with STAR v2.7.1a^21^ according to ENCODE protocol^22^. DEseq2 DE analysis was then conducted with the same parameters as described above for mRNAs.

#### Deconvolution of RNA-seq data

To account for potential tissue heterogeneity, we used a murine atrial gene signature matrix described in Donovan et al.^23^ and our matrix of gene expression in Fragments Per Kilobase of exon model per Million reads mapped (FPKM) in CIBERSORTx online tool^24^. We then performed nonparametric Wilcoxon test on all possible comparisons for fibroblast fraction with a statistical significance threshold of *p* < 0.05.

#### Gene set enrichment analyses

For each gene sets described above, we performed hypergeometric testing against the human Gene Ontology (GO) Biological Processes (BP) from Molecular Signatures Database v7.1 with the HypeR package.

#### miRNA target prediction

For DE miRNA present in the 5 most cited miRNA databases (DIANA, Miranda, PicTar, TargetScan, and miRDB), we defined genes as targets if they were: i-annotated with a human homolog in the ensemble database, ii-predicted targets by at least 3 out of 5 databases queried with the MiRNAtap package, iii-DE (mRNA FDR < 0.01), iv-inversely correlated (Pearson’s *r*<−0.5) log_2_ expression, corrected for the fibroblast fraction (expression values corrected with Limma’s removeBatchEffect() function). We then performed a GSEA with the remaining 82 predicted targets of the miRNA located on the syntenic region of the Dlk1-Dio3 locus (CanFam3.1 Chr8:68961744-69696779) as described above.

#### RNA-seq and miRNA-seq DE genes comparison between human AF patients and canine AF models

DE genes in our canine AF models with annotated human orthologues in the ENSEMBL database were compared to a meta-analysis of miRNA DE in human AF and a large RNAseq study on left atrial appendages obtained from 261 patients undergoing valve surgery^25, 26^ Precursors of the human miRNAs listed in Shen et al. Table S8 (*n= 53, 21 upregulated and 32 downregulated*)^25^ were retrieved with the R package miRBaseConverter and then compared across species. Because only 5 miRNA were found to overlap with human DE miRNA, only mRNA genes found to be DE in human and our canine AF models are represented as an Upset plot. The counts for human mRNA data were downloaded from GEO database (GSE69890). DE testing was conducted as described above for the 3 groups; no AF (CTL, n=50), AF in AF rhythm (AF, n=130), and AF in sinus rhythm (AF.SR, n=81), with inclusion of sex as a covariate.

#### Mitochondrial genes DE in canine AF models

The human MitoCarta3.0^27^ database was queried for genes with mitochondrial localization in the heart (n=539). We represented DE genes with likelihood ratio test FDR<0.01 from that list as volcano plots.

### Proteomics

Dog cardiomyocytes were lysed by sonication, reduced and alkylated. Protein was precipitated, resuspended, quantified and subjected to tryptic digest. Peptides (500 ng) were analyzed by reverse phase nano-HPLC coupled to a Bruker maXis II mass spectrometer (positive mode, mass range 150 - 2200 m/z, collision induced dissociation of top 20 precursors). LC-MS/MS data were analyzed for protein identification and label-free quantification using MaxQuant^28^ (1.6.1.0) against the public database UniProt with taxonomy Canis lupus familiaris and common contaminants (downloaded on 01.08.2019, 29809 sequences) with carbamidomethylation on Cys as fixed and oxidation on Met as variable modification with decoy database search included (mass tolerance 0.006 Da for precursor, 80 ppm for product ions; 1 % PSM and protein FDR, match between runs enabled, minimum of 2 ratio counts of quantified razor and unique peptides).

#### DE analysis and correlation

Proteins with > 3 missing values per treatment were removed. The remaining missing intensities were replaced with random values taken from the Gaussian distribution centered around a minimal value from the 10th quantile with the *DEP* package’s Minprob function, to simulate a relative label-free quantification (LFQ) value for those low abundant proteins. Two-sample t-tests with subsequent multiple testing correction by FDR were used to identify DE proteins (*p*<0.01) with the fibroblast fraction as covariate using the Limma package.

Because proteomic processing does not always converge to a single protein, only 755 genes out of the 1029 in the proteomic matrix were correlated to their corresponding RNA-seq data. We compared overlapping genes’ mean log_2_ transformed expression in proteomic and RNA-seq. The distribution of mean RNA-seq expression of the 755 overlapping genes was then compared to the full mean RNA-seq gene expression values.

## RESULTS

### RNA-sequencing of cardiomyocyte-enriched atrial samples from canine AF models

We analyzed data from three groups of six dogs. The first group (CTL) was the control group without atrioventricular ablation (AVB) nor pacemaker, in the second group (AF), right atrial-tachypacing at 600 beats per minute (b.p.m.) was used to maintain AF electrically for one week, and the third group (AF+AVB) included dogs with electrically-maintained AF for one week in the presence of AVB and ventricular pacing at 80 b.p.m. to control the ventricular rate. We reasoned that transcriptomic profiling of atria from these animals should allow us to discover the molecular changes that occur over the first week after the onset of AF, and play a role in the development of the tissue remodeling accompanying the transition from paroxysmal to persistent AF.

Initial analysis of bulk RNA-seq data hinted at some heterogeneity of cellular composition across samples. Therefore, we estimated the fraction of the major cell types in each sample using an *in-silico* deconvolution technique implemented in CIBERSORTx (Fig. 1A)^24^. Because of the induced tissue remodeling due to the AF treatments, we found that both AF and AF+AVB dogs had more fibroblasts in their atria than CTL animals (Fig. 1B). To emphasize the transcriptional differences between conditions that are not a result of variable cellular composition, we included the fibroblast fraction as a covariate in all subsequent DE analyses. Correction for this confounding variable reduced inter-group variability (Fig. 1C-D).

**Figure 1.**
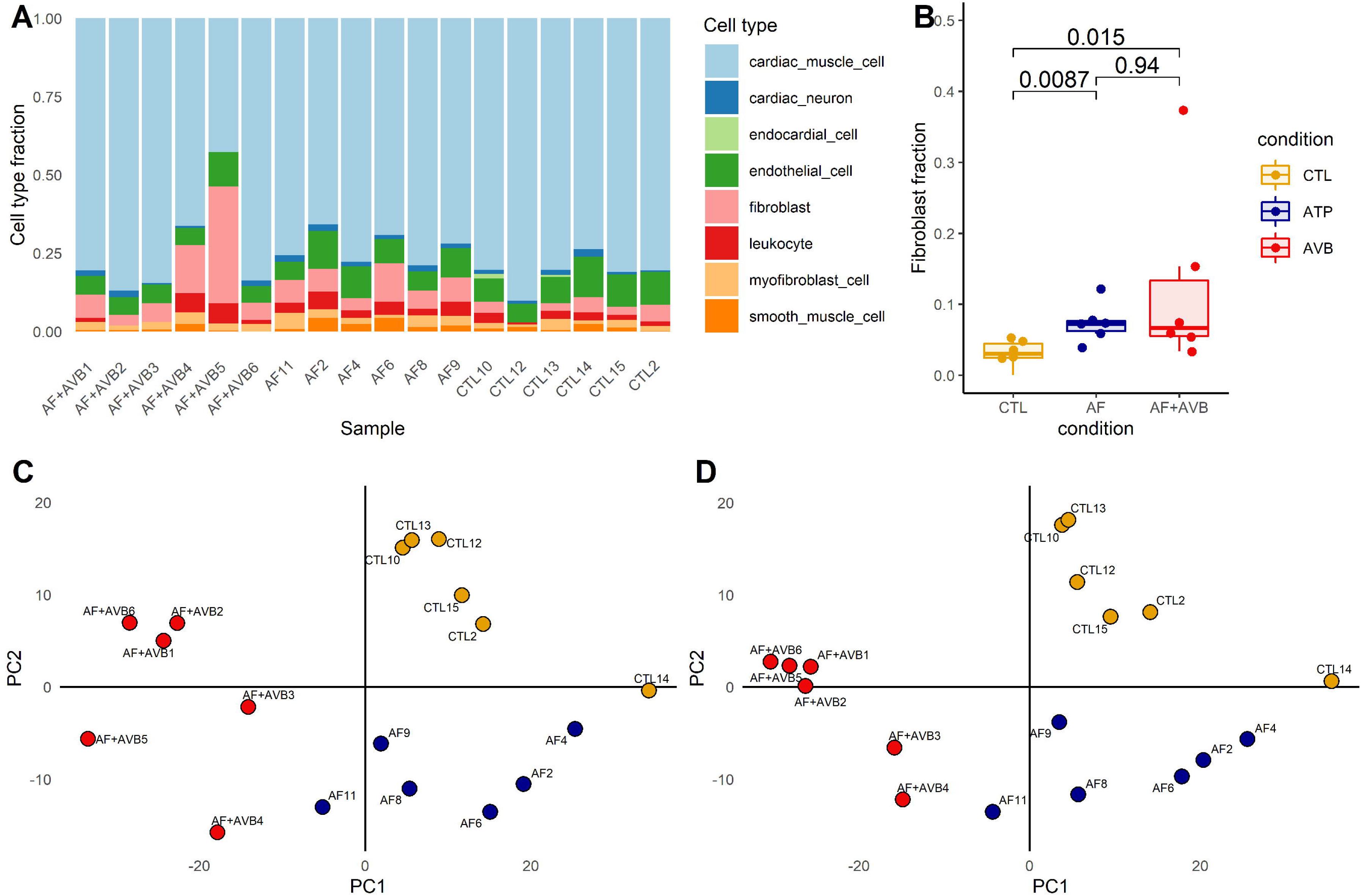
Deconvolution of canine atria cell composition using bulk RNA-sequencing. (**A**) We inferred cell fractions with CIBERSORTx and an atrial-specific gene signature matrix obtained using orthologous murine genes^23^. We present cell fractions for each dog sample that we analyzed in this study. CTL, control; AF, Atrial-tachypacing; AF+AVB, AF with Atrio-Ventricular Block. (**B**) When we group animals per treatment arm, we observed a significantly higher fraction of fibroblasts in the atrial fibrillation dog models (AF and AF+AVB) than in the control animals (AFvsCTL Wilcoxon’s test *P*=0.0087 and AF+AVBvsCTL *P*=0.015). Principal component analysis of the top 1000 most variable genes expressed in canine atria before **(C)** and after **(D)** correction for fibroblast fraction show treatment-dependent clustering after correction for cell composition.

### Proteomic analysis largely confirms the transcriptomic results

To validate our RNA-seq results, we took advantage of mass spectrometry (MS)-based protein quantification results from the same 18 dog atrial cardiomyocyte-enriched cell extractions that were generated in a parallel study (detailed proteomic results will be presented elsewhere). After stringent quality control, we obtained relative quantification for 755 proteins. For these genes, the relative RNA and protein levels were strongly correlated (Pearson’s *r*=0.49, *P*=1.57×10^-46^) (**Fig. 2A**). Many of the genes that are well-correlated encode abundant cardiomyocyte proteins, such as titin (*TTN*), myosin light chain-4 (*MYL4*), desmin (*DES*), and tropomyosin-1 (*TPM1*). We found that RNA-seq could profile transcripts with a wider range of expression profiles, whereas MS-based proteomics preferentially captured proteins whose genes are expressed at high levels. (**Fig. 2B**).

**Figure 2.**
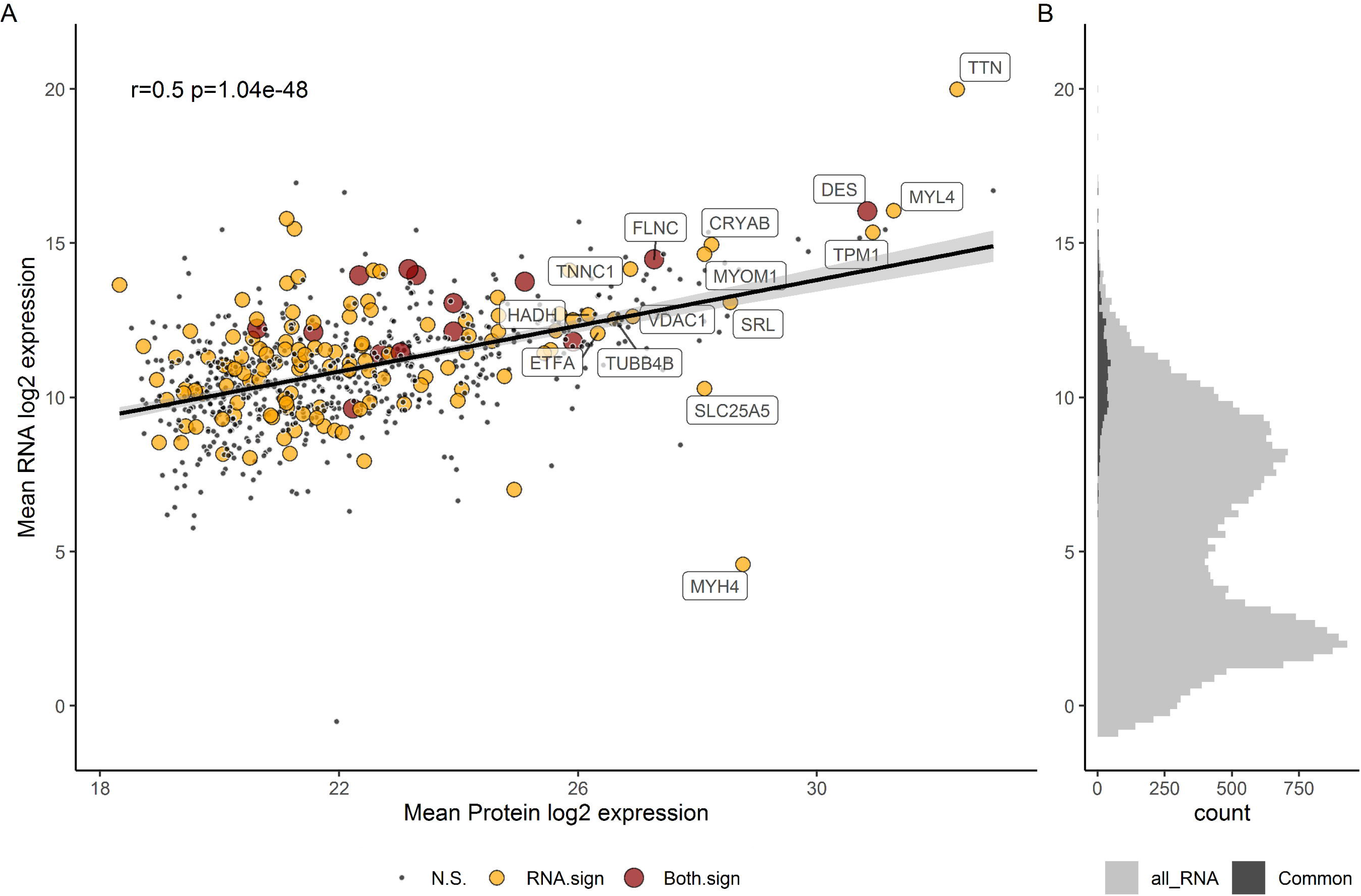
Validation of highly expressed RNA by proteomics. (**A**) In 18 atrial samples, 755 genes (N.S.= 619, RNA.sign=122, Both.sign=14) found in both datasets are highly correlated at the protein (x-axis) and RNA (y-axis) levels (Pearson’s *r*=0.49, *P*=1.57×10^-46^). For reference, we annotated 15 genes that are differentially expressed in the RNA-seq experiment and have high protein expression levels. N.S., not differentially expressed in the RNA-seq or proteomic experiment; RNA.sign, genes that are differentially expressed in the RNA-seq assay only; Both.sign, differentially expressed genes in both the RNA-seq and proteomic experiments. DE genes in the RNAseq dataset have an FDR < 0.01 (likelihood ratio test) and proteomics dataset an FDR < 0.05 (F-test). The grey area around the line corresponds to the 95% confidence interval. (**B**) Relative expression level of all transcripts measured in the RNA-seq experiment. The histogram shows that genes that are present in both the RNA-seq and proteomic experiment are highly expressed (Common, dark grey) in comparison to the expression levels of all transcripts measured (all_RNA, light grey).

### Transcriptomic changes in cardiomyocyte-enriched atrial samples

Pairwise comparisons of gene expression levels between the three groups of dogs identified 434, 5971, and 7867 genes that are DE (false discovery rate (FDR) <0.01) in atrial cardiomyocyte-rich fractions in AFvsCTL, AF+AVBvsCTL, and AFvsAF+AVB, respectively (**Fig. 3A-B**). All differential gene expression level results are available in **Supplemental Table II** and https://github.com/lebf3/DogAF). Many genes previously implicated in AF are dysregulated in both AF and AF+AVB dogs when compared to controls, thus validating the experimental design. This includes *FHL1* involved in myofilament regulation^29^, *SORBS2* involved in intercalated disc gap junction regulation^30^, and *KCNA5*, which regulates atrial action potential repolarization^31^. Previous studies have established an important role for mitochondrial dysfunction in the etiology of AF^27^. Accordingly, we identified 54 genes that encode mitochondrial proteins that are DE in our AF canine models (**Supplemental Figure I**).

**Figure 3.**
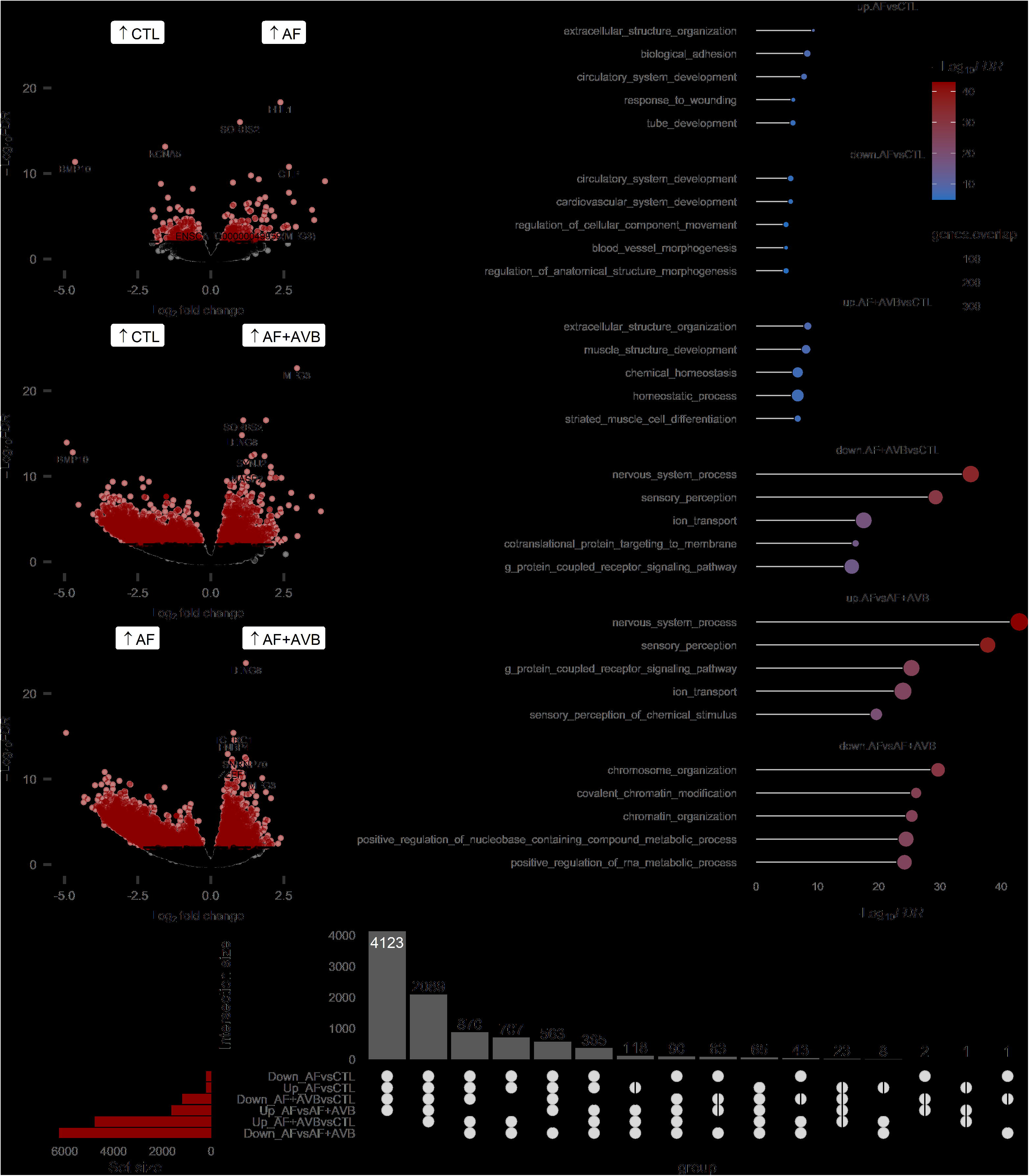
Analyses of differentially expressed atrial genes identify many biological pathways that are dysregulated in atrial fibrillation dog models. (**A**) Volcano plots of all transcripts that we analyzed in this study. Transcripts in red have a false discovery rate (FDR)<0.01. We found 434, 5971 and 7867 genes that were DE in the AFvsCTL, AF+AVBvsCTL, and AFvsAF+AVB analyses, respectively. The full DE results are available in **Supplemental Table II**. (**B**) Upset plot showing the intersection of up -and down-regulated DE genes (FDR<0.01) in each analysis. (**C**) The five most significant biological pathways identified using gene-set enrichment analyses (GSEA) for each set of DE genes (FDR <0.01). Full results are available in **Supplemental Table IV**.

In particular in the AF+AVB group, we noted the up-regulation of two key beta-oxidation genes (*CPT1A* and *ACADL*) and the down-regulation of the electron transport chain genes *COX17* and *NDUFA8*. However, our data also implicates genes not previously recognized to be involved in AF, such as leukocyte receptor cluster member-8 (*LENG8*), transcription elongation regulator-1 (*TCERG1*), ligand dependent nuclear receptor corepressor (*LCOR*), formin-binding protein-4 (*FNBP4*), and *ENSCAFG00000049959* (orthologue of the lncRNA *MEG3*)(**Fig. 3A**, **Supplemental Table III**, and https://github.com/lebf3/DogAF).

To understand what pathways are modulated in the atria of these canine AF models, we performed gene set enrichment analyses (GSEAs) on the DE genes (**Fig. 3C** and **Supplemental Table IV**). In AFvsCTL, we noted an up-regulation of genes associated with profibrotic pathways (*e.g*. extracellular structure organization, biological adhesion, response to wounding) and a down-regulation of genes implicated in angiogenesis, such as blood vessel morphogenesis. Genes implicated in muscle biology were up-regulated in the AF+AVBvsCTL analysis (*e.g*. muscle structure development, striated muscle cell differentiation) whereas the same comparison implicated down-regulated genes involved in ion transport and signaling pathways (*e.g*. sensory perception). We confirmed that this enrichment was not due to a smaller fraction of cardiac neurons found in the atria of AF+AVB dogs (Kruskal-Wallis’ *P*=0.32). Because of the large overlap in genes that are down-regulated in AF+AVBvsCTL and up-regulated in AFvsAF+AVB (**Fig. 3B**), we identified similar pathways in the GSEA for these two comparisons (in **Fig. 3C**, compare AF+AVBvsCTL and AFvsAF+AVB). Finally, genes that were down-regulated in the AFvsAF+AVB analysis implicated genes with more generic functions in gene expression and chromatin modifications, such as the histone-lysine N-methyltransferase *SETD5* and the DNA methyltransferase *TET2*. Dysregulation of the expression of these chromatin-related genes and pathways is consistent with the extensive transcriptomic changes observed in the atria of AF+AVB dogs, in sheep models of AF (**Supplemental Figure II**) as well as in AF patients^32, 33^.

### Dysregulation of miRNA expression

Because miRNA play important roles in AF biology^34^ but are not detected in standard RNA-seq protocols, we performed in parallel miRNA-seq on the same dog samples. We found 31, 19 and 21 miRNA that are DE (FDR <0.01) in AFvsCTL, AF+AVBvsCTL and AFvsAF+AVB, respectively (**Fig. 4A**, **Supplemental Table II)**. When comparing miRNA expression in the two AF models, *MIR185* on the dog chromosome 26 was the most DE miRNA with strong upregulation in the atria of AF animals. We also noted that 11 of the most strongly DE miRNA in the AF+AVBvsCTL and AFvsCTL analyses (*MIR136, MIR411, MIR370, MIR127, MIR493, MIR494, MIR485, ENSCAFG00000025655* (96.20% identity to hsa-mir-379), *MIR758, MIR543, MIR889*) mapped to the chr8:68,900,000-69,700,000 region in the dog reference genome CanFam3.1 (**Fig. 4B**). This region, highly conserved in mammals, is syntenic to the imprinted 14q32 region in humans (also known as the *DLK1-DIO3* locus)^35^. The lncRNA *MEG3*, which we described above as being over-expressed in the AF canine models is also located in the same *DLK1-DIO3* syntenic dog locus.

**Figure 4.**
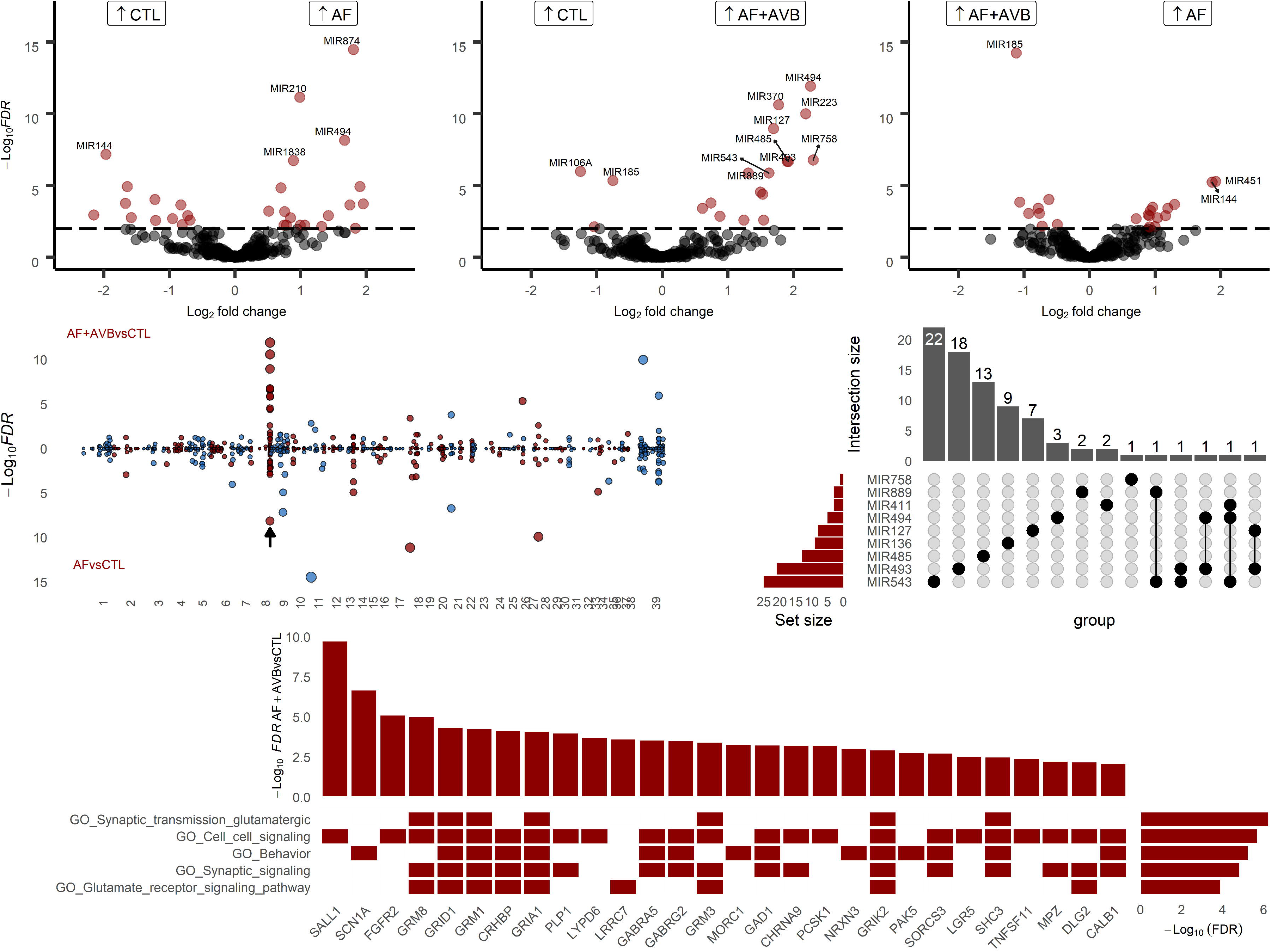
11 differentially expressed microRNA*s* (miRNA*s*) map to a canine chromosome 8 region that is syntenic to human DLK1-DIO3. (**A**) Volcano plots of all miRNA that we measured in our experiments. We identified 31, 19 and 20 miRNA that are differentially expressed (false discovery rate (FDR) <0.01) in the AFvsCTL, AF+AVBvsCTL and AFvsAF+AVB analyses, respectively. (**B**) Miami plots of miRNA and their corresponding statistical significance (y-axis) for the AF+AVBvsCTL (top) and ATvsCTL (bottom) analyses. An arrow indicates the miRNA cluster located on the canine chromosome 8 region that is syntenic to human DLK1-DIO3. The odd and even chromosomes FDR values are in blue and red respectively. (**C**) Upset plot showing the DE miRNA targets located in the syntenic DLK1-DIO3locus and their corresponding number of potential target RNA. We identified potential targets with the MiRNAtap package (predicted by ≥ 3 databases) from DE miRNA (FDR <0.01) and DE mRNA (FDR<0.01). (**D**) Gene-set enrichment analyses (GSEA) with the potential gene targets (x-axis) of the DE miRNA located at the syntenic DLK1-DIO3 locus. We only present the top five pathways enriched in this analysis. A red square in the heatmap indicates membership of a given target gene to the biological pathways located on the left (empty columns were removed for clarity). GSEA FDR and AF+AVBvsCTL DE FDR are on the right and top of the heatmap, respectively.

The dysregulation of the expression of lncRNA and miRNA at the same locus suggested that they might co-regulate the expression of genes implicated in the same biological pathway(s). To address this possibility, we used in silico predictions to infer the DE mRNA that are possible direct targets of these DE miRNA located at the syntenic *DLK1-DIO3* locus. For this analysis, we focused on DE miRNAs and DE mRNAs that are predicted to physically interact by at least three out of five databases and that have expression levels that are negatively correlated in the RNA-seq/miRNA-seq experiments (Pearson’s *r* < −0.5). Using these filters, we identified 82 potential target genes for the DE miRNAs at this locus, with most genes targeted by a single miRNA (**Fig. 4C**). GSEA with these 82 genes indicated a common role in synaptic signaling involving glutamate signaling (**Fig. 4D** and **Supplemental Table V**). Some of the key genes within these pathways are metabotropic glutamate receptor-1 and −8 (*GRM1, GRM8*), glutamate ionotropic receptor delta type subunit-1 (*GRID1*), glutamate ionotropic receptor AMPA type subunit-1 (*GRIA1*), and corticotropin-releasing factor-binding protein (*CRHBP*).

### Partial differential transcriptomic overlap between human and dog AF atrial samples

To assess the ability of our canine AF models to capture early transcriptomic changes which might be missed by profiling the atria of human AF patients who have developed the disease over years, we compared the DE genes identified in AF dogs with results from human left atrial transcriptomic profiling experiments^25, 26^ Hsu et al. performed bulk RNA-seq experiments on left atrial appendages from 261 patients who underwent cardiac surgery to treat AF, valve disease, or other cardiac disorders. For differential gene expression analyses, these patients were divided between no AF (n=50), AF in AF rhythm (n=130), and AF in sinus rhythm (n=81). When we intersected this list of human DE genes with the list of DE genes in our AF dog models (with clear human orthologs), we identified 668 genes (Fig. 5 and Supplemental Table VI). We found the strongest overlap between dog AF+AVB and human AF in AF rhythm. Of note, most of the strongest signals in our dog study are also present in this human study (*LENG8, SORBS2, BMP10, FNBP4* and glutamate receptor-related genes (*GRM1, GRM8, GRIA1, GRIK2, GRID1)*). For miRNA, we compared our results with data from a large meta-analysis involving 40 articles and 283 DE miRNA in AF (in different tissues and species)^25^. Of the 53 AF-associated miRNAs that were previously identified in human cardiac tissues and showed consistent results in the meta-analysis, we found five miRNAs in our analyses of the dog transcriptomic datasets (*MIR144, MIR142, MIR146B, MIR223* and *MIR451*). These miRNAs have not been characterized functionally yet for a role in AF. Generally, the canine AF group matched better the directionality of change reported in the human AF DE miRNA meta-analysis.

**Figure 5.**
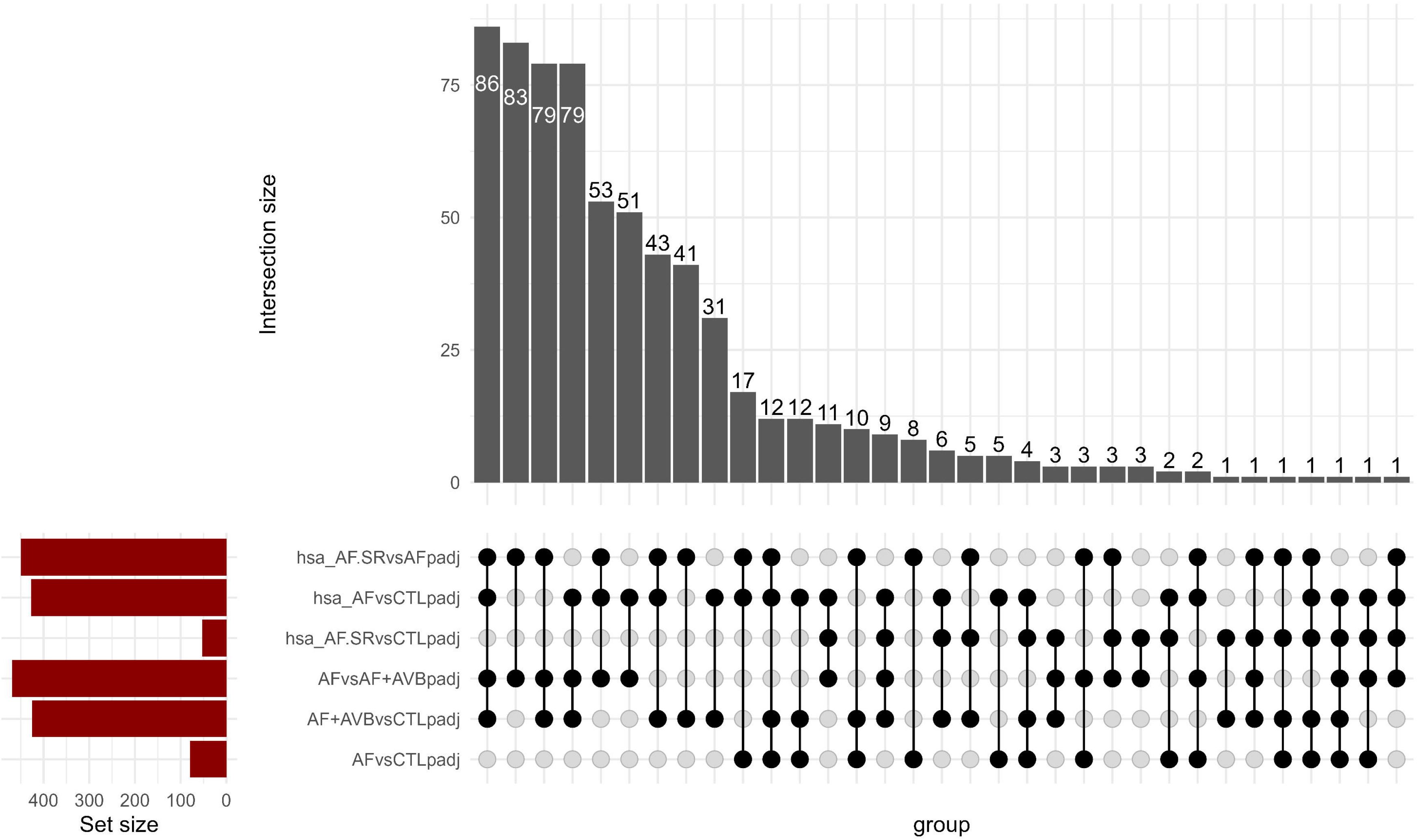
Overlaps in genes differentially expressed in canine AF models and human AF patients. We compared differentially expressed genes in our canine AF models with annotated human orthologues that are DE in human AF left atrial appendages^26^. Homo Sapiens; hsa, hsa_AF; AF in AF rhythm, hsa_AF.SR; AF in sinus rhythm, hsa_CTL; no AF.

## DISCUSSION

In this study, we used a transcriptomic approach to comprehensively assess the molecular architecture of AF-induced remodeling with and without AVB from atria cardiomyocyte-enriched samples. We validated the robustness of our RNAseq data by correlating it with a proteomic analysis, which showed a strong correlation in gene expression, with well-defined cardiomyocyte genes being most highly expressed in both datasets (e.g. TTN, MYL4). We confirmed the involvement of known AF factors like the reactivation of developmental pathways, but also found a strong and novel association with microRNAs and lncRNA from the *DLK1-DIO3* locus, including the *MEG3* canine orthologue. This finding is concordant with the many chromatin remodeling genes dysregulated in our models, which is an emerging phenotype of AF both in human and sheep models^36^.

### Molecular remodeling in AF with versus without AVB

We did not expect to find a smaller number of DE genes in the AFvsCTL analysis, given our previous observation that AF treatment alone without AVB results in more important tissue remodeling^10^. One possible explanation is that our prior histological studies were done in AF animals treated for three weeks,^10^ whereas the results presented here reflect RNA changes after one week of AF. The transcriptomic changes in the AF+AVB group show that cells are under active chromatin modification, indicating ongoing adaptation to the stimulus. This is not observed in the AF group (lacking AVB), which may indicate that this adaptation has already occurred. This idea would be consistent with the down-regulation of chromatin-related genes recently noted in the atria of sheep AF models^36^, and may be a result of earlier establishment of profibrotic transdifferentiation in AF compared to AF+AVB canine models.

Our analyses highlighted many genes not previously implicated in AF. While the functions of some of these genes remain uncharacterized (*e.g. LENG8* in AF+AVB), we can speculate on the activities of others. For instance, *TCERG1* and *FNBP4*, which are up-regulated in the atria of AF+AVB dogs, encode co-regulated proteins that are involved, respectively, in RNA splicing and translation^37^. The up-regulation of these genes, with general actions on gene transcripts, may (partly) explain why more genes are dysregulated in AF+AVB animals when compared to the CTL or AF groups (**Fig. 3B**). Another interesting candidate is *LCOR*, which is up-regulated in the AF+AVB model and encodes a transcriptional cofactor that interacts with PPARγ and RXRα to control gene expression^38^. While further experiments are needed to determine the extent by which *LCOR* modifies gene expression in AF+AVB and contributes to the pathology, our RNA-seq experiments detected the dysregulation of two of its likely targets based on the literature: *CPT1A* (see above) and the cell cycle regulator *CDKN1A*^39^.

### Potential role of non-coding genes at the *DLK1-DIO3* locus in early AF

The highly-conserved *DLK1-DIO3* locus hosts two differentially DNA-methylated regions modulating the expression of its non-coding RNA clusters, where in humans the maternal allele is hypomethylated with concomitant expression on the hypermethylated paternal allele of non-coding RNA and other protein-coding genes (*DLK1, RLT1*, and *DIO3*)^35^. In both AF+AVBvsCTL and AFvsCTL, we found a large proportion (58% and 23%, respectively) of DE miRNA at this locus, underlying its importance in AF-related adaptation. We also found dysregulation of the *MEG3* lncRNA canine orthologue at this locus. *MEG3* is a highly expressed lncRNA that has been studied in various pathologies, including cancer^40^ and more recently cardiovascular diseases^41^. Non-coding RNAs at this locus have been shown to mediate various cardiac developmental programs^35^. More specifically, *MEG3* can contribute to the recruitment of the Polycomb repressive complex-2 (*PRC2*)^42^, a key chromatin modulating factor. Of particular interest, Mondal et *al*. showed that through interaction with the H3-Lys-27 methyltransferase *EZH2, MEG3* can repress TGF-beta target genes, which are known to promote a profibrotic response^43^. Data have been presented that suggest an important role of *EZH2* and/or *EZH2*-regulated genes in AF^44^.

### Glutamate receptor regulation by miRNAs from the *DLK1-DIO3* locus

Our GSEA analysis-predicted gene targets of DE miRNA at the *DLK1-DIO3* locus suggest a role for glutamate signaling in AF. Immunostaining has confirmed the presence of glutamate receptors on cardiomyocytes^45^. Glutamate was also found to be significantly increased in AF patient left atrial appendages^46^. Glutamate signaling is important in vagal afferent neurons^47^, and remodeling of the glutamate system in AF may relate to the extensive previous evidence of autonomic dysfunction in AF patients^48^. Moreover, a recent study has shown fundamental roles for glutamatergic receptors in rat atrial cardiomyocytes and induced pluripotent stem cell-derived atrial cardiomyocytes, including a reduction in cardiomyocyte excitability after *GRIA3* knockdown^49^. Therefore, the *DLK1-DIO3* miRNA cluster may be an adaptative regulator of cardiomyocyte excitability or of neural cells in the presence of AF.

### Limitations

We used cardiomyocyte-enriched samples in an attempt to obtain clearer results from the transcriptomic analysis by excluding extrinsic variability due to changes in cell composition. However, while our samples are enriched in cardiomyocytes, they do not constitute a pure cardiomyocyte preparation. A disadvantage is that variability due to changes in cell composition is not eliminated. On the other hand, our cardiomyocyte-enriched (but not pure) samples allow us to detect potential features of AF related to non-cardiomyocyte cells, such as autonomic dysregulation mediated by neural cells; however, we cannot unambiguously attribute DE genes to transcriptomic changes in a specific cell type. In part, we were able to control for fibroblast composition by adjustment through analysis for expression of fibroblast-related RNA-expression patterns. Nevertheless, features underlined here should be confirmed in pure cell cultures or single cell transcriptomic assays. A second limitation is the difficulty in extrapolating our findings to gene expression changes in humans. We found only modest overlap of DE genes in our model compared to reported DE gene patterns in human; several factors could explain this (*e.g*. differences in biospecimen preparation, tissue heterogeneity, fundamental differences between dog and human AF pathology). It is also possible that different transcriptomic programs may be involved at the initiation of arrhythmia and tissue remodeling (AF and AF+AVB dog models) when compared with those dysregulated in the atria once the pathology has been present for years.

We compared our transcriptomic results with proteomic data on the same samples and found a very high level of correlation (**Fig. 2**). These results are consistent with a large measure of transcriptomic control over protein expression and validate the relevance of transcriptomic analysis of these data. A further in-depth look at the proteomic signature in these models would be of interest but is beyond the scope of the present manuscript.

### Conclusions

Understanding the pathophysiology of chronic human diseases such as AF is challenging because they develop over many years and initially present with only unremarkable pre-clinical symptoms. In this study, we took advantage of two well-characterized canine AF models to chart the transcriptomic changes that occur at the earlier phases of arrhythmia.

Despite the inherent limitations in relating dog models to human AF, our results offer interesting new hypotheses for future testing, including in man. In particular, the up-regulation of miRNAs at the *DLK1-DIO3* locus after 1 week of AF suggests that they may be early biomarkers of tissue remodeling and/or adaptation in the atria.

## Nonstandard abbreviations and acronyms

CTL: Control
AF+AVB: Atrial tachypacing with atrioventricular-node ablation
AF: Atrial tachypacing without atrioventricular-node ablation
RAA: Right atrial appendage
LA: Left atrium
DE: Differential expression
GSEA: Gene set enrichment analysis
lncRNA: Long non-coding RNA

## Supplemental Materials

Supplementary Table I & Supplementary Figures I-II

Supplementary Tables II-VI

---

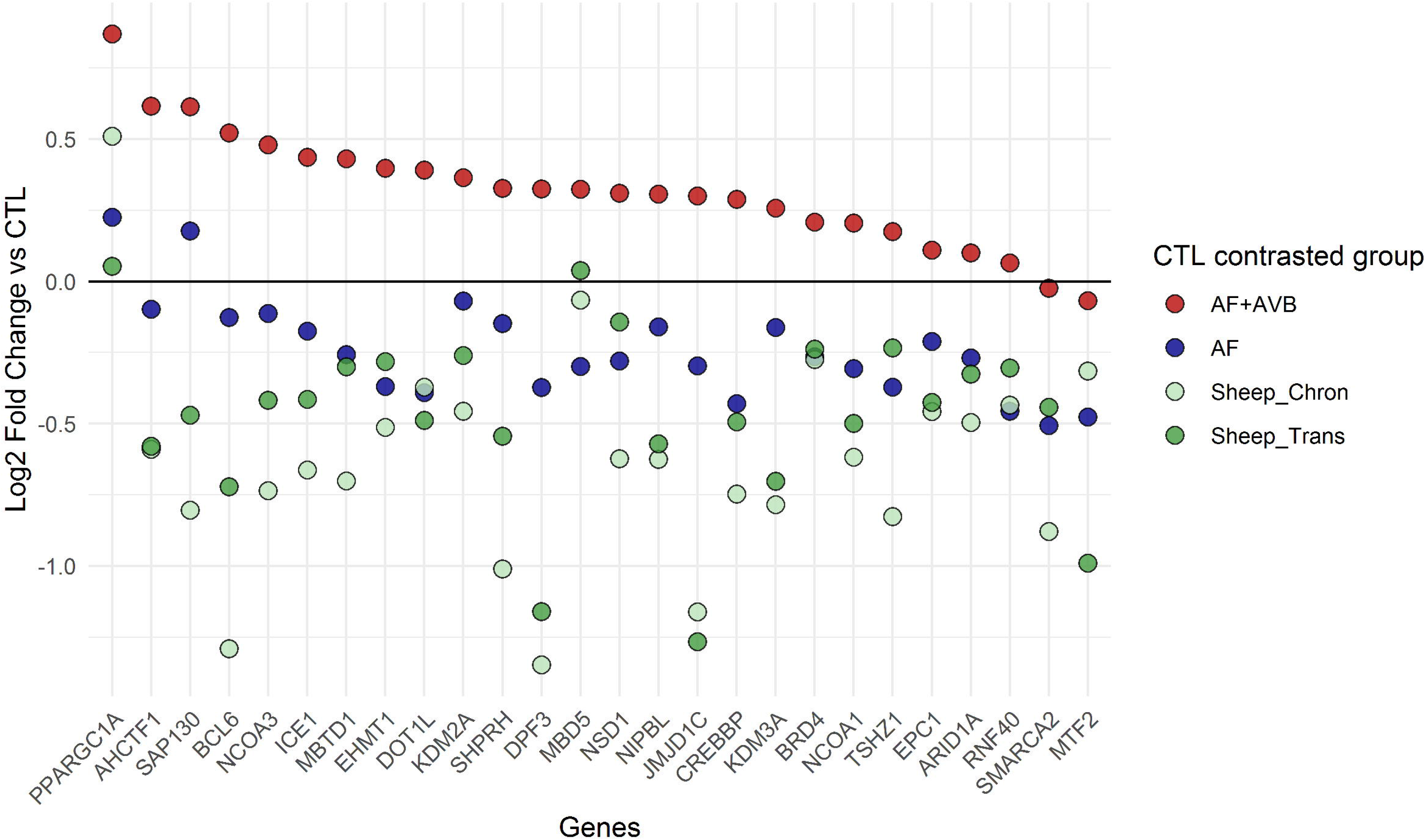

